# Strand Displacement Chain Reaction (SDCR): New Hybrid Amplification Technique for Fast and Sensitive Detection of Genetic Materials

**DOI:** 10.1101/2025.07.31.667828

**Authors:** Evgeniya V. Smirnova, Ekaterina V. Barsova, Dmitriy A. Varlamov, Vladimir M. Kramarov, Konstantin A. Blagodatskikh, Konstantin B. Ignatov

**Author notes:** **Correspondence:** Konstantin B. Ignatov, Vavilov Institute of General Genetics, Russian Academy of Sciences, 119991 Moscow, Russia. (E.V.S.); (E.V.B.).

## Abstract

Nucleic acid amplification methods are widely used in science, medicine and forensics for molecular biological assays and for the detection of genetic material. The newly developed strand displacement chain reaction (SDCR) method is a hybrid amplification technique based on polymerase chain reaction (PCR) and isothermal nucleic acid amplification. Here, we compared conventional PCR, the “gold standard” for molecular diagnostic assays, with the SDCR method by performing real-time amplification assays using human, bacterial and viral genetic materials. In the assays, SDCR demonstrated very high sensitivity and amplification efficiency. We found that the SDCR method provided an amplification factor above three, which noticeably outperformed that of PCR amplification and enabled a marked reduction in the number of cycles in comparison with PCR. Therefore, the new hybrid amplification technique could be extremely useful for the detection of genetic material and the development of new diagnostic kits.

## 1. Introduction

Nucleic acid (NA) amplification methods are widely used in biotechnology, molecular biology, genetics, forensics and medicine for molecular diagnostic assays. Currently, two main strategies are used to amplify defined sequences of nucleic acids: thermocycling DNA amplification by polymerase chain reaction (PCR) and isothermal amplification.

The “gold standard” of sequence-specific DNA amplification is polymerase chain reaction (PCR), with it being the oldest and most widely used [1–3]. This reaction relies upon instrument-based thermal cycling on the denaturation of a template DNA, followed by the annealing of primers at specific sites on the denatured template and their extension by a thermostable DNA polymerase (such as Taq DNA polymerase [4–6]), allowing for an exponential increase in the amount of DNA [2]. The main advantages of this method include its high sensitivity, the specificity of the reaction and the prevalence of the required equipment.

Isothermal amplification methods emerged in the early 1990s and gradually gained popularity thanks to the simple equipment required and the speed of the process in isothermal mode. Though these methods may require an initial high-temperature incubation to denature the template DNA for the initiation of the process, DNA amplification takes place at one defined temperature, while the separation of the template DNA strands during amplification is carried out by enzymatic strand displacement activity. A variety of isothermal amplification methods have been developed, including the following: nucleic acid sequence-based amplification (NASBA) [7], strand displacement amplification (SDA) [8,9], rolling circle amplification (RCA) [10], cross priming amplification (CPA) [11], nickase-dependent amplification (NDA) [12] and loop-mediated amplification (LAMP) [13]. These methods, like many other isothermal amplification methods, require the use of a DNA polymerase with strong strand displacement activity [14,15], such as Phi29 [16] or Bst DNA polymerase [15,17]. Generally, isothermal amplification provides faster assay rates than PCR amplification, but PCR provides a higher assay sensitivity [18].

Upon the creation of DNA polymerases that possess both high thermal stability and high strand displacement activity, such as SD DNA polymerase [14], it became possible to combine the processes of thermocyclic and isothermal amplification in one reaction. This has led to the emergence of a number of new hybrid methods that combine the advantages of isothermal amplification and PCR. Among the hybrid NA amplification methods, those of particular interest include polymerase chain displacement reaction (PCDR) [14,19], coupled PCR-LAMP [18] and pulse-RCA [20,21]. PCDR is a hybrid technique combining PCR with strand displacement amplification; coupled PCR-LAMP combines a PCR-like thermocycling mode at the initial stage of amplification with an isothermal LAMP mode during the following stage; and pulse-RCA is a combination of rolling circle amplification with PCR. The hybrid methods advantageously combine both the high sensitivity of PCR and the fast assay rate of isothermal amplification [18].

Recently, a new method of hybrid DNA amplification was developed by Dong et al., named strand displacement chain reaction (SDCR) [22]; the reaction scheme of this method is shown in Figure 1. This method appears as a hybrid of nickase-dependent isothermal amplification and PCR. SDCR requires deoxyribooligonucleotide primers that flank the DNA fragment to be amplified, similar to primers used for PCR, and contain one (or more) ribonucleotides. Said ribonucleotides should be located within the oligonucleotide sequences of the primers, dividing the primers into 3’ and 5’ terminal parts. In addition to the primers, the reaction mixture should contain a thermostable DNA polymerase with a strong strand displacement activity (SD DNA polymerase) and a thermostable ribonuclease H type II (RNase H2), which acts as a nickase in the reaction. SDCR amplification of the target DNA sequence is performed under a thermal cycling mode, for which the reaction mixture is successively heated to the denaturation temperature of dsDNA and then cooled to the temperature at which primer annealing and the enzymatic synthesis of new DNA chains occurs. During the annealing and elongation of primers on the template DNA, thermostable RNase H2 introduces a single-strand break (nick) in the nucleotide sequence of the annealed primers immediately upstream of the ribonucleotide(s) contained in the primers. The resulting nicks serve as the initiation sites for the synthesis of a new DNA chain, and the chain initially synthesized as a result of primer elongation is displaced by the chain-displacing activity of DNA polymerase. The result is a double-stranded DNA amplification product and displaced single-stranded DNA. The displaced DNA strand is primed by a complementary oligonucleotide primer containing a ribonucleotide(s), which again leads to primer elongation, the introduction of a break in the nucleotide sequence of the primer, and the displacement of the new DNA strand with the simultaneous formation of a double-stranded amplification product. Thus, at the primer annealing and elongation step, the process of DNA synthesis is continuous and performed in an isothermal mode. This ensures a more than twofold increase in the copy number of the amplified DNA template in one thermal cycle. It should be noted that the double-stranded DNA amplification product is not involved in isothermal amplification. In order to involve double-stranded DNA in the amplification process, a thermal denaturation step is required, resulting in double-stranded DNA (dsDNA) being converted into single-stranded DNA (ssDNA), which serves as a template for the synthesis of new DNA strands.

**Figure 1.**
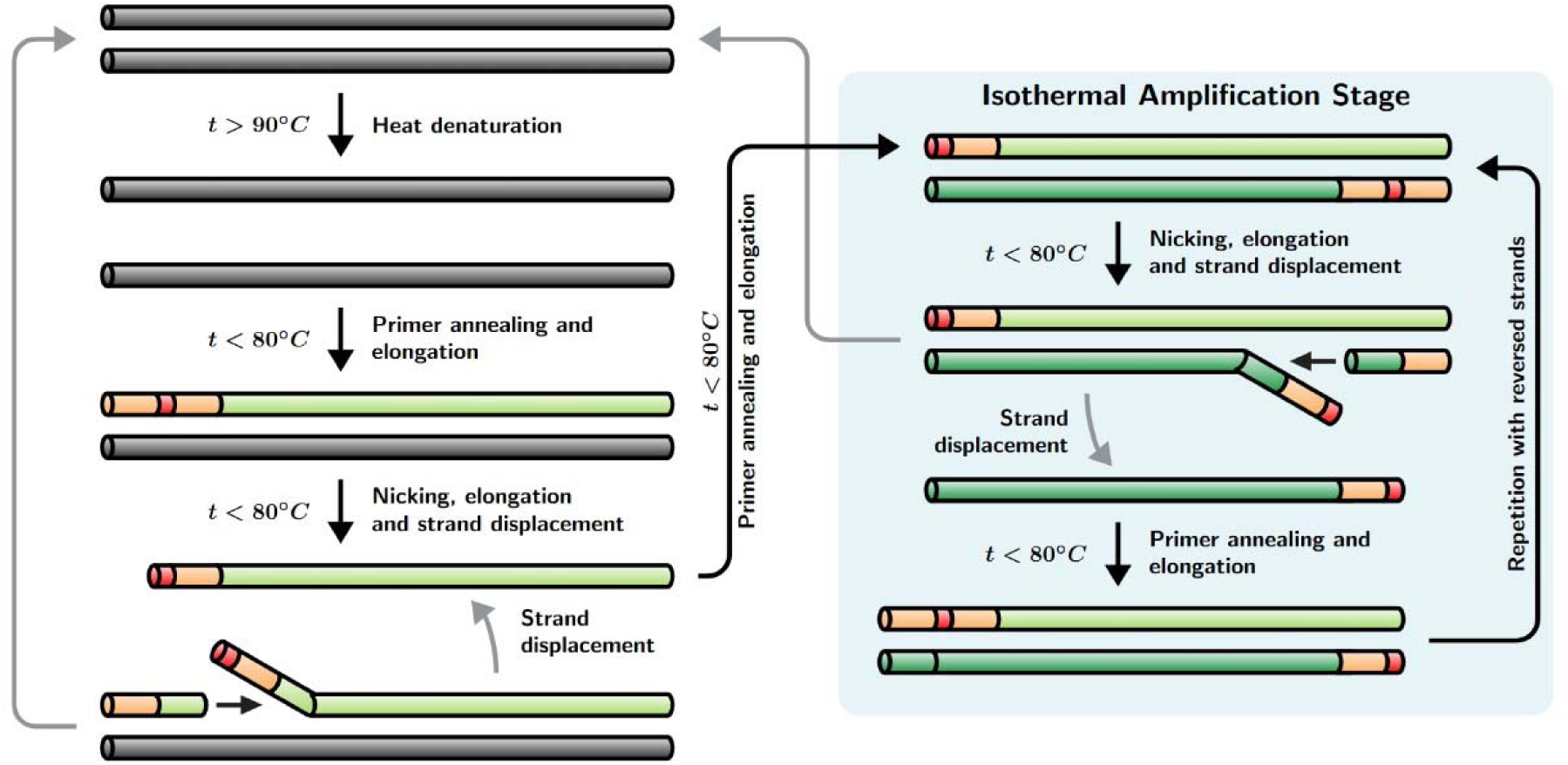
Schematic diagram of SDCR amplification. A description of the amplification process is given in Section 1. The initial template DNA is shown as dark gray pipes. Primers are shown as peach-colored, and ribonucleotides are indicated in red. Newly synthesized DNA strands are shown as green pipes (direct strands in light green and reverse strands in dark green). The presence of the isothermal amplification stage enables increased template amplification per thermocycle compared with standard PCR.

The potential advantages of SDCR include a remarkably higher amplification efficiency than that of of PCR, the ability to use standard PCR analysis equipment, and the ability to perform the reaction in “real-time” mode using a hydrolysis probe [22]. The use of hydrolysis probes capable of complementary interactions with the amplified sequences enables a significant increase in the specificity of the analysis. To perform SDCR with a hydrolysis probe, the latter should contain a ribonucleotide in its sequence to ensure its hydrolysis by RNase H2 after hybridization with the target sequence.

Conceivably, the use of SDCR could greatly improve amplification assays, but there have not yet been any examples of practical applications of the technique, except for amplifications from plasmid templates [22]. In the present work, we evaluate the performance of quantitative SDCR (qSDCR) and compare it with quantitative PCR (qPCR) in the real-time amplification assays of target sequences from human cDNA and bacterial and viral gDNA.

## 2. Materials and Methods

### 2.1. Enzymes and Reagents

The following materials were obtained from their respective sources: SD HotStart DNA polymerase (10 U/µL), SmartHot Taq DNA polymerase (5 U/µL), and 10× SD polymerase reaction buffer from Bioron GmbH (Römerberg, Germany); dNTPs from Bioline Limited (London, UK); oligonucleotide primers and probes were synthesized by Syntol JSC (Moscow, Russia); the cDNA library of human embryonic kidney 293 (HEK 293) cells and viral gDNA of lambda bacteriophage from Evrogen JSC (Moscow, Russia); and human gDNA and bacterial gDNA of *Mycobacterium bovis* (stain Russia BCG-1) from Syntol JSC. Concentrations of the used DNA samples were verified using the Quant-iT PicoGreen dsDNA Assay Kit (Invitrogen by ThermoFisher Scientific, Inc., USA).

Thermostable RNase H2 (Pa RNase H2) was isolated from a recombinant *E. coli* strain that carries the RNase H2 gene from *Pyrococcus abyssi.* The cloning, expression, and purification of recombinant Pa RNase H2 are fully described in [23]. The enzyme is also commercially available from Integrated DNA Technologies, Inc. (Coralville, Iowa, USA).

The real-time amplification reactions were performed using a CFX96 Touch Real-Time Detection System (Bio-Rad Laboratories, Inc., Hercules, CA, USA).

### 2.2. cDNA of RNA isolated from FFPE blocks

Samples of total RNA were isolated from two separate formalin-fixed, paraffin-embedded (FFPE) blocks containing fixed human tissues using the RNeasy FFPE Kit for RNA Extraction (Qiagen GmbH, Hilden, Germany). Total RNA concentration was measured with a Qubit RNA High-Sensitivity kit and a Qubit-4 fluorimeter (ThermoFisher Scientific, Inc., Carlsbad, CA, USA).

cDNA was obtained from 500 ng of each RNA sample by reverse transcription using the HiScript III 1st Strand cDNA Synthesis Kit (Vazyme, Nanjing, China) according to the protocol with DNase I treatment in a volume of 20 μL. A specific primer to beta-2-microglobulin (B2M) RNA (5’-CAATCCAAATGCGGCATCTTC) was used for priming the 1st strand of cDNA. The synthesized cDNA was used in the following assays without additional purification.

### 2.3. Real-time qSDCR amplification

qSDCR assays were performed in real-time mode with a hydrolysis oligonucleotide probe and modified primers for SDCR, all of which contained ribonucleotide in the middle of their sequences. The 3’ terminal parts of the SDCR primers and the corresponding conventional PCR primers were completely identical and complementary to the amplified sequences, but the SDCR primers additionally contained artificially generated 5’-end sequences upstream of the ribonucleotide (the actual sequences are provided in Sections 2.5-2.7). The reactions were carried out with an enzyme mixture of SD HotStart DNA polymerase and Pa RNase H2. The SD polymerase has no nuclease activity, so the reactions with this DNA polymerase contained Pa RNase H2 for the probe hydrolysis and for the nicking of SDCR primers. The reaction mixtures (25 µL) contained 0.25 mM dNTP (each); two modified primers comprising a ribonucleotide (0.2 µM, each); a hydrolysis oligonucleotide probe containing a ribonucleotide (0.3 µM); 3 mM MgCl_2_; a specified amount of a template DNA, or a no-template control (NTC); SD HotStart DNA polymerase (20 U); Pa RNase H2 (2.5 ng); and 1X SD-reaction buffer. Preliminarily, we optimized the concentration of the SD polymerase and found that concentrations over 0.8 U/µL did not improve the performance of SDCR (Figure S1).

The following conditions were used for SDCR performance: (a) initial denaturation at 92 °C for 90 s; and (b) amplification via 40 thermocycles at 92 °C for 30 s, and 66 °C for 60 s.

### 2.4. Real-time qPCR amplification

qPCR assays were performed using a hydrolysis oligonucleotide probe, conventional PCR primers, and Taq polymerase or an enzyme mixture of SD polymerase and Pa RNase H2. The SD polymerase has no nuclease activity, so RNase H2 was needed for the probe hydrolysis. The reaction mixtures (25 µL) contained 0.25 mM dNTP (each); two non-modified primers (0.2 µM, each); a hydrolysis oligonucleotide probe containing a ribonucleotide (0.3 µM); 3 mM MgCl_2_; a specified amount of a template DNA, or a no-template control (NTC); SmartHot Taq DNA polymerase (5 U), or an enzyme mixture of SD HotStart DNA polymerase (20 U) with Pa RNase H2 (2.5 ng); and 1X SD-reaction buffer. Preliminarily, we optimized concentrations of the enzymes used in the reactions. We found that Taq polymerase concentrations over 0.2 U/µL or SD polymerase concentrations over 0.4 U/µL did not improve the performance of PCR [14].

### 2.5. qSDCR and qPCR assays of lambda bacteriophage gDNA

The real-time SDCR assay was performed as described in Section 2.3 with the following probe and primers: *PL*-probe (5’(ROX)-CCGGCG(rG)CTGCAGGT(BHQ2)GCAGAGTG-(P)), and primers *PL-f1* (5’- GTGCCGTGAAGACGACAGAGTT(rG)GTGGTCTGCCCTGGCTGAGTGA) and *PL-r1* (5’- GCAGTGGTATCTCGCAGGAGTT(rG)GCAGACCAGCCACCACGGCA).

The PCR detection of lambda gDNA (GenBank Accession NC_001416) was performed as described in Section 2.4 with the following probe and primers: *PL*-probe (above); primers *PL-f2* (5’-GGTGGTCTGCCCTGGCTGAGTGA) and *PL-r2* (5’-GGCAGACCAGCCACCACGGCA).

The size of the target amplified sequence was 76 bp. The reactions were performed with a series of 10-fold dilutions of the input template, from 250 fg to 0.25 fg of lambda gDNA per reaction. All reactions (including a no-template control, NTC) contained 50 ng of human gDNA.

### 2.6. *qSDCR and qPCR assays of* Mycobacterium bovis gDNA

The real-time assays were performed as described in Sections 2.3 and 2.4. For the detection of *M. bovis* gDNA (GenBank Accession NZ_CP013741) by SDCR amplification, the following probe and primers were used: *MT-*probe (5’-(ROX)-GCGGATCTCT(rG)CGACCAT(BHQ2)CCGCACCGCCCGC-(P)), and primers *MT-f1* (5’- GTGCCGTGACGACGACAGCGTT(rG)CTGCCCACTCCGAATCGTGCT) and *MT-r1* (5’- GCAGTGGTCTC TCGCAGGCGTT(rG)TACCCGCCGGAGCTGCGT).

For the PCR assays. we used *MT-*probe (above) and primers *MT-f2* (5’- GCTGCCCACTCCGAATCGTGCT) and *MT-r2* (5’-GTACCCGCCGGAGCTGCGT).

The size of the target amplified sequence was 78 bp. The reactions were carried out with a series of 10-fold dilutions of the input template, from 11 pg to 11 fg of *M. bovis* gDNA per reaction. Also, all reactions (including a no-template control, NTC) contained 50 ng of human gDNA.

### 2.7. qSDCR and qPCR assays of human B2M cDNA

The real-time SDCR and PCR assays of human beta-2-microglobulin (B2M) cDNA (GenBank Accession BC_064910) were performed as described in Sections 2.3 and 2.4. The human cDNA library or B2M cDNA synthesized from RNA isolated from FFPE blocks (Section 2.2) was used as a template. The reactions using the cDNA library were carried out with a series of 10-fold dilutions of the input template, from 10 pg to 0.01 pg of the library per reaction. The reactions with the B2M cDNA of RNA isolated from FFPE blocks were performed with 1 μL of the non-purified cDNA.

The following probe and primers were used for the SDCR assays: *B2M-*probe (5’-(ROX)-CACAGCCCAA(rG)ATAGTT(BHQ2)AAGTGGGATCGAGAC-(P)), and primers *B2M-f1* (5’- GTGCCGTGAAGACGACAGAGTT(rG)CCTGCCGTGTGAACCATGTGACT) and *B2M-r1* (5’- GCAGTGGTATCTCGCAGGAGTT(rG)CCAAATGCGGCATCTTCAAACCTCCA).

The *B2M-*probe (above) and the following primers were used for the PCR assays: *B2M-f2* (5’- GCCTGCCGTGTGAACCATGTGACT) and *B2M-r2* (5’- CCAAATGCGGCATCTTCAAACCTCCA).

The size of the B2M amplified sequence was 101 bp.

### 2.8. Data Analysis

All data processing was performed using R software [24]. Core R language and several extension packages were used, including RDML for raw amplification data manipulation [25], chipPCR for Cq calculation [26], and ggplot2 for the generation of plots [27].

The efficiency of each amplification variant was estimated by standard curve (dilution series of template DNA). The log of each concentration in the dilution series (x-axis) was plotted against the Cq value for that concentration (y-axis). Then, efficiency was determined by the following equation, using the slope of the standard curve: Efficiency = 10^(D1/slope)^ [1. Amplification per one cycle (amplification factor) was determined as 10^(D1/slope)^.

## 3. Results

### 3.1. qSDCR and qPCR assays of lambda bacteriophage gDNA

The sensitivity and efficiency of the qSDCR and qPCR techniques were evaluated in detecting phage lambda gDNA. The tests were carried out as described in Section 2.5 and we used a series of 10-fold dilutions of the target DNA, from 250 fg to 0.25 fg of lambda gDNA per reaction. Conventional qPCR assay with Taq polymerase was used as a standard. Additionally, qPCR and qSDCR tests were performed with a mixture of SD DNA polymerase and Pa RNase H2. Figure 2 and Table 1 show the results of the assays.

**Figure 2.**
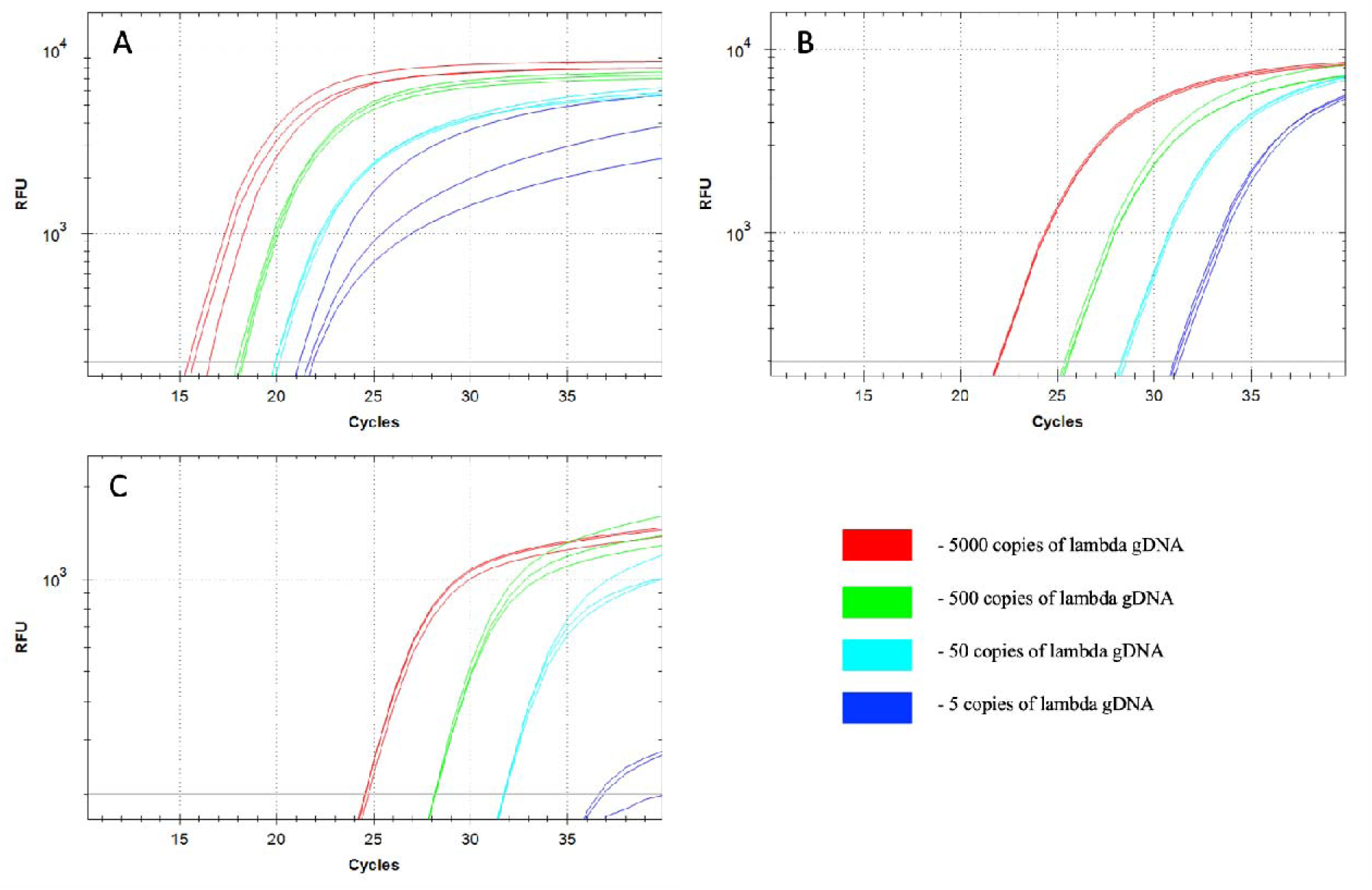
Comparison of qSDCR and qPCR assays of *Lambdavirus lambda* gDNA. The quantitative real-time reaction assays contained 5000, 500, 50, or 5 copies of lambda bacteriophage gDNA per reaction, or a no-template control (NTC). The NTC samples did not produce a detectable fluorescent signal. The reactions (including a no-template controls) contained 50 ng of human gDNA. The detection of viral DNA was performed by qSDCR (A) and qPCR (B, C) assays. The PCR assays were carried out with SD (B) and Taq (C) DNA polymerases. qSDCR markedly outperformed qPCR assays.

**Table 1.** Comparison of quantitative SDCR and PCR assays of phage λ gDNA. The mean quantification cycles (Cq) ± standard deviation of three replicates for the indicated amounts of the bacterial gDNA (NTC, no-template control) are provided. ΔCq1 is the difference between the Cq of the qSDCR and the qPCR with SD polymerase (qPCR-1), and ΔCq2 is the difference between the Cq of the qSDCR and the control qPCR with Taq polymerase (qPCR-2). The efficiency of amplification variants is expressed by the slope of the standard curve, the PCR efficiency, and the amplification factor.

| **Template (amount of phage $\lambda$ gDNA) per reaction** | **qSDCR**<br>**(SD pol + RNase H2)** | **qPCR-1**<br>**(SD pol + RNase H2)** | **qPCR-2**<br>**(Taq pol)** |
| --- | --- | --- | --- |
| 250 fg<br>(5000 copies) | Cq 15.79 ± 0.55<br>ΔCq1 □ 6.16<br>ΔCq2 □ 8.76 | Cq 21.95 ± 0.06 | Cq 24.55 ± 0.1 |
| 25 fg<br>(500 copies) | Cq 18.09 ± 0.12<br>ΔCq1 □ 7.31<br>ΔCq2 □ 10.03 | Cq 25.4 ± 0.1 | Cq 28.12 ± 0.02 |
| 2.5 fg<br>(50 copies) | Cq 19.97 ± 0.12<br>ΔCq1 □ 8.35<br>ΔCq2 □ 11.72 | Cq 28.32 ± 0.1 | Cq 31.69 ± 0.04 |
| 0.25 fg<br>(5 copies) | Cq 21.52 ± 0.37<br>ΔCq1 □ 9.58<br>ΔCq2 □ 15.19 | Cq 31.1 ± 0.11 | (Cq 36.71 ± 0.17) <sup>2</sup> |
| NTC | N/A <sup>1</sup> | N/A <sup>1</sup> | N/A <sup>1</sup> |
| **Efficiency of amplification** |  |  |  |
| Slope | -1.91 | -3.04 | -3.57 |
| qPCR efficiency | 234 % | 113 % | 91 % |
| Amplification factor | 3.34 | 2.13 | 1.91 |

<sup>1</sup> N/A: Not available.
<sup>2</sup> This mean Cq value was calculated by the use of two replicates (the third replicate had failed and was not considered for the efficiency of qPCR with Taq polymerase).

Standard qPCR with Taq polymerase demonstrated a lower amplification efficiency than the qPCR with SD polymerase, which was seen as a 2–3 cycle lag in the assays. This fact is in agreement with earlier described findings [14].

The qSDCR significantly outperformed the qPCR tests and yielded much better Cq values ([ΔCq approximately 8–15 cycles for different template dilutions). As the result, the qSDCR assay reduced the number of thermocycles by a third in comparison with the qPCR. The efficiency of amplification can be estimated by the amplification factor (amplification per one cycle). It was 3.34 for qSDCR and approximately 2 (1.91-2.13) for qPCRs (Table 1).

Although both qSDCR and qPCR methods had high sensitivity levels and allowed the detection of up to 5 copies of viral genomic DNA per reaction, qSDCR allowed us to detect this amount at 21–22 cycles, whereas qPCRs detected it only at 31–37 cycles, demonstrating the significant advantage of qSDCR (Table 1). Although all reactions (including no-template controls) contained 50 ng of human gDNA, the assays did not generate false-positive results.

### 3.2. qSDCR and qPCR assays of Mycobacterium bovis gDNA

The qSDCR and qPCR assays of the bacterial DNA were performed similarly to the assays of the viral DNA described above. A series of 10-fold dilutions of *M. bovis* gDNA, from 11 pg to 11 fg, was used. The reactions (including no-template controls) additionally contained 50 ng of human gDNA and further details of the assays are described in Section 2.6. For the detection of *M. bovis,* insertion sequence IS6110 was selected, as it is a targeting sequence commonly used to detect the Mycobacterium tuberculosis complex (MBTC) [28]. The genome of the *M. bovis* stain Russia BCG-1 contains two copies of IS6110.

Figure 3 and Table 2 show the results of the detection tests. The qPCR assays with Taq and SD polymerases demonstrated similar results. The qSDCR again significantly outperformed the qPCRs and reduced Cq values by more than a third ([ΔCq approximately 11–17 cycles for different dilutions of the template). The amplification factor for qSDCR was 3.2 and 1.91-1.95 for qPCR (Table 2). Again, although both assays provided high sensitivity and could detect as few as 2.5 copies of bacterial genomic DNA or 5 copies of the target sequence per reaction, the number of cycles for detection was 35–36 for PCR, compared to only 19–20 needed for the SDCR.

**Figure 3.**
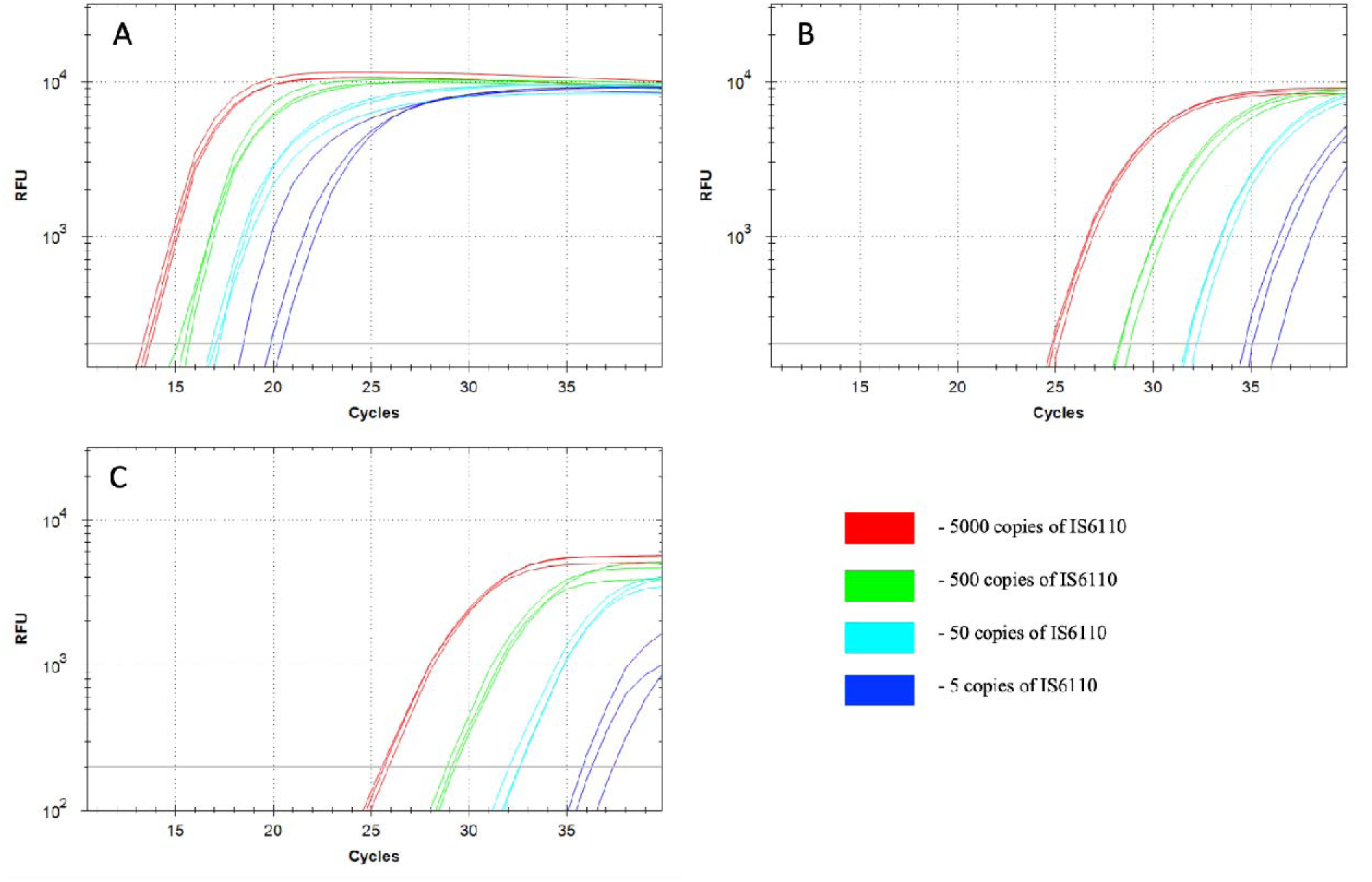
Comparison of qSDCR and qPCR assays of *Mycobacterium bovis* gDNA. The quantitative real-time reaction assays contained 5000, 500, 50, or 5 copies of the target IS6110 sequence per reaction, or a no-template control (NTC); the NTC samples did not produce a detectable fluorescent signal. The reactions (including no-template controls) contained 50 ng of human gDNA. The detection of *M. bovis* gDNA was performed by qSDCR (A) and qPCR (B, C) assays, while PCR assays were carried out with SD (B) and Taq (C) DNA polymerases. qSDCR significantly outperformed qPCR assays.

**Table 2.** Comparison of quantitative SDCR and PCR assays of *M. bovis* gDNA. The mean quantification cycles (Cq) ± standard deviation of three replicates for the indicated amounts of the bacterial gDNA (NTC, no-template control) are provided. ΔCq1 is the difference between the Cq of the qSDCR and the qPCR with SD polymerase (qPCR-1); and ΔCq2 is the difference between the Cq of the qSDCR and the control qPCR with Taq polymerase (qPCR-2). The efficiency of amplification variants is expressed by the slope of the standard curve, the PCR efficiency, and the amplification factor.

| **Template (amount of *M. bovis* gDNA) per reaction** | **qSDCR**<br>**(SD pol + RNase H2)** | **qPCR-1**<br>**(SD pol + RNase H2)** | **qPCR-2**<br>**(Taq pol)** |
| --- | --- | --- | --- |
| 11 pg<br>(2500 genome copies) | Cq 13.4 ± 0.19<br>ΔCq1 □ 11.48<br>ΔCq2 □ 12.21 | Cq 24.88 ± 0.2 | Cq 25.61 ± 0.2 |
| 1.1 pg<br>(250 genome copies) | Cq 15.29 ± 0.23<br>ΔCq1 □ 13.06<br>ΔCq2 □ 13.74 | Cq 28.35 ± 0.3 | Cq 29.03 ± 0.2 |
| 0.11 pg<br>(25 genome copies) | Cq 16.97 ± 0.2<br>ΔCq1 □ 14.8<br>ΔCq2 □ 15.35 | Cq 31.77 ± 0.31 | Cq 32.32 ± 0.26 |
| 0.011 pg<br>(2.5 genome copies) | Cq 19.45 ± 1.04<br>ΔCq1 □ 15.81<br>ΔCq2 □ 16.94 | Cq 35.26 ± 0.87 | Cq 36.39 ± 0.8 |
| NTC | N/A <sup>1</sup> | N/A <sup>1</sup> | N/A <sup>1</sup> |
| **Efficiency of amplification** |  |  |  |
| Slope | -1.98 | -3.46 | -3.56 |
| qPCR efficiency | 220 % | 95 % | 91 % |
| Amplification factor | 3.2 | 1.95 | 1.91 |

<sup>1</sup> N/A: Not available.

### 3.3. qSDCR and qPCR assays of B2M cDNA from human cDNA library

We compared the efficiency of the qSDCR with the qPCR technique in the detection tests of human beta-2-microglobulin (B2M) cDNA. The assays were carried out as described in Section 2.7. A series of 10-fold dilutions of the human total cDNA library, from 10 pg to 10 fg of the library per reaction, was used for the testing. The results are shown in Table 3 and Figure 4.

**Figure 4.**
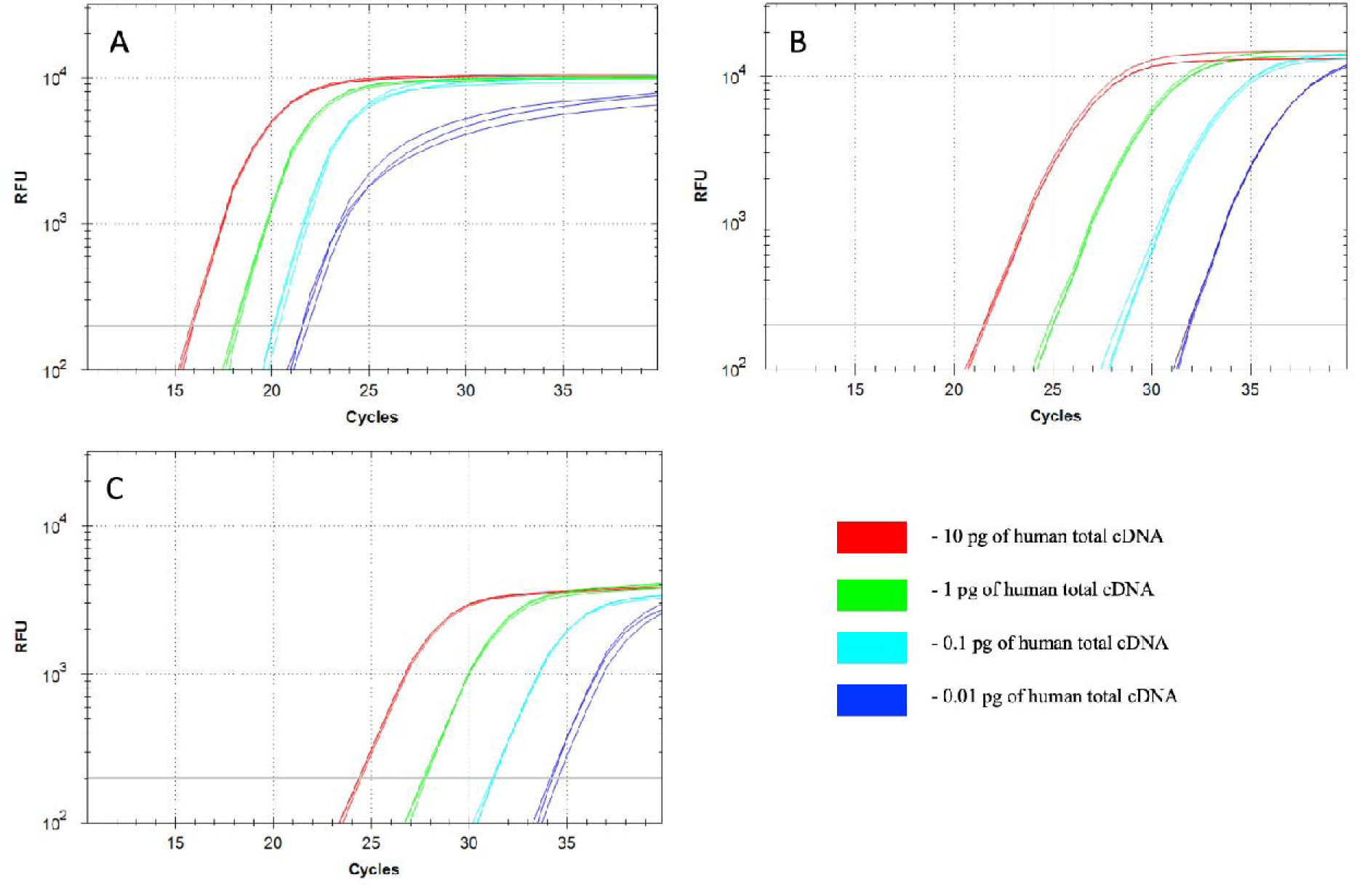
Comparison of qSDCR and qPCR assays of human B2M in cDNA library. The reaction assays contained 10, 1, 0.1, or 0.01 pg of the human total cDNA library per reaction, or a no-template control (NTC). The NTC samples did not produce a detectable fluorescent signal. The assays were performed by qSDCR (A) and qPCR (B, C). The PCR tests were carried out with SD (B) and Taq (C) DNA polymerases. qSDCR noticeably outperformed qPCR assays.

**Table 3.**
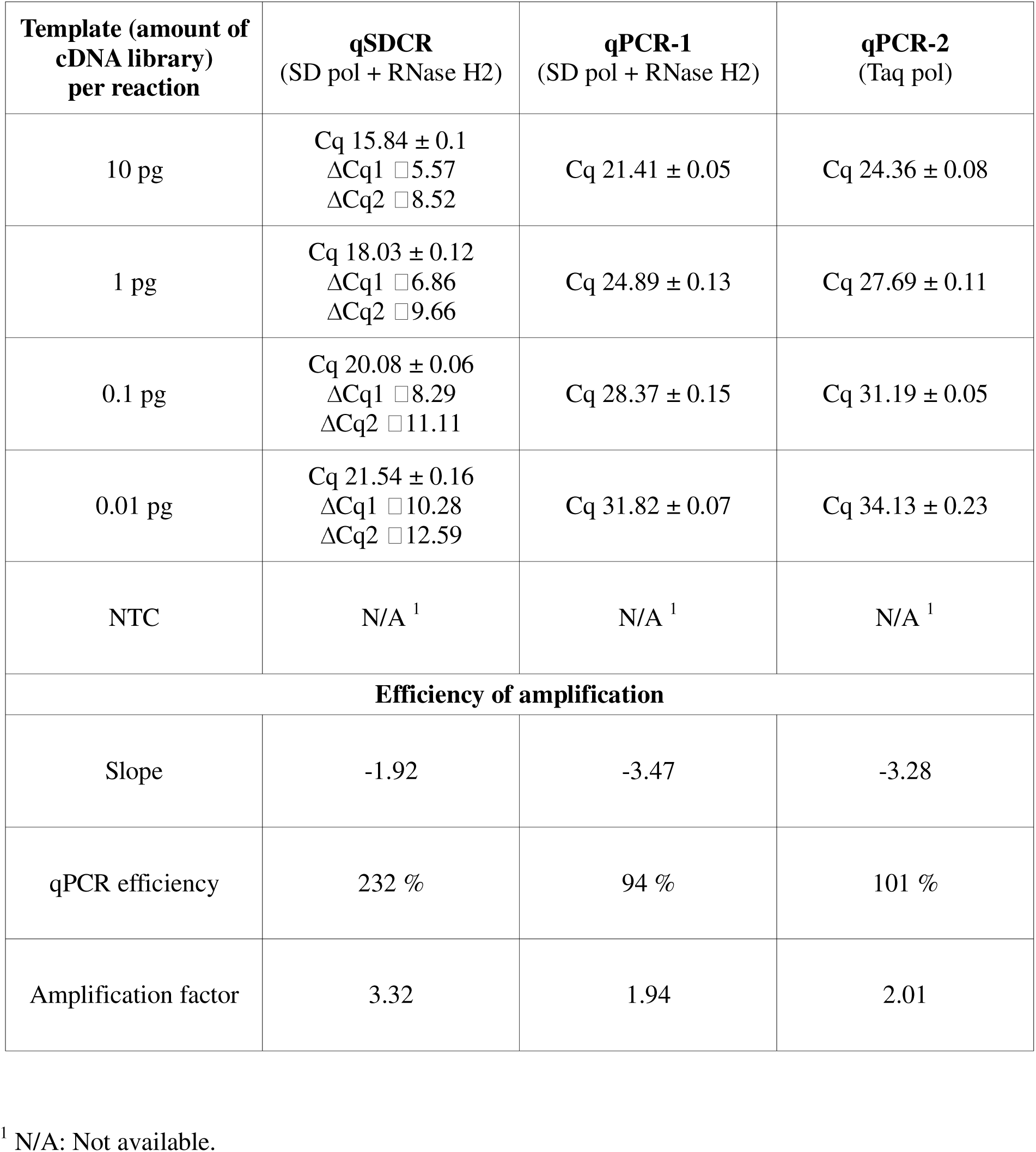
Comparison of quantitative SDCR and PCR assays of human B2M cDNA. The mean quantification cycles (Cq) ± standard deviation of three replicates for the indicated template total cDNA amounts (NTC, no-template control) are provided. ΔCq1 is the difference between the Cq of the qSDCR and the qPCR with SD polymerase (qPCR-1); and ΔCq2 is the difference between the Cq of the qSDCR and the control qPCR with Taq polymerase (qPCR-2). The efficiency of amplification variants is expressed by the slope of the standard curve, the PCR efficiency, and the amplification factor.

| **Template (amount of cDNA library) per reaction** | **qSDCR**<br>**(SD pol + RNase H2)** | **qPCR-1**<br>**(SD pol + RNase H2)** | **qPCR-2**<br>**(Taq pol)** |
| --- | --- | --- | --- |
| 10 pg | Cq 15.84 ± 0.1<br>ΔCq1 □ 5.57<br>ΔCq2 □ 8.52 | Cq 21.41 ± 0.05 | Cq 24.36 ± 0.08 |
| 1 pg | Cq 18.03 ± 0.12<br>ΔCq1 □ 6.86<br>ΔCq2 □ 9.66 | Cq 24.89 ± 0.13 | Cq 27.69 ± 0.11 |
| 0.1 pg | Cq 20.08 ± 0.06<br>ΔCq1 □ 8.29<br>ΔCq2 □ 11.11 | Cq 28.37 ± 0.15 | Cq 31.19 ± 0.05 |
| 0.01 pg | Cq 21.54 ± 0.16<br>ΔCq1 □ 10.28<br>ΔCq2 □ 12.59 | Cq 31.82 ± 0.07 | Cq 34.13 ± 0.23 |
| NTC | N/A <sup>1</sup> | N/A <sup>1</sup> | N/A <sup>1</sup> |
| **Efficiency of amplification** |  |  |  |
| Slope | -1.92 | -3.47 | -3.28 |
| qPCR efficiency | 232 % | 94 % | 101 % |
| Amplification factor | 3.32 | 1.94 | 2.01 |

<sup>1</sup> N/A: Not available.

The SD polymerase provided a better qPCR performance than that of Taq polymerase ([ΔCq approximately 2-3 cycles). The qSDCR dramatically outperformed the qPCR and provided much better Cq values than those with the qPCR technique with Taq or SD DNA polymerase ([ΔCq approximately 6–12 cycles for different dilutions of the template). The amplification factor was 3.32 for qSDCR and 1.94-2.01 for qPCR tests.

### 3.4. qSDCR and qPCR assays of human B2M cDNA obtained from FFPE tissue blocks

The efficiency of the qSDCR and qPCR techniques in detecting genetic material obtained from FFPE tissue blocks was evaluated by testing the cDNA of human beta-2-microglobulin (B2M). Two samples of B2M cDNA were obtained from total RNA isolated from two separate formalin-fixed, paraffin-embedded (FFPE) blocks containing human tissues as described in Section 2.2. The assays were carried out with the non-purified cDNA as described in Section 2.7. Figure 5 demonstrates that qSDCR surpasses qPCR in the efficiency of the amplification assays. The Cq values of the qSDCR tests were 19 cycles for sample 1 and 21.2 cycles for sample 2, while those for the qPCR tests were 27.4 cycles for sample 1 and 29.9 cycles for sample 2. Thus, the qSDCR tests reduced the number of thermocycles by approximately a third in comparison with the qPCR tests.

**Figure 5.**
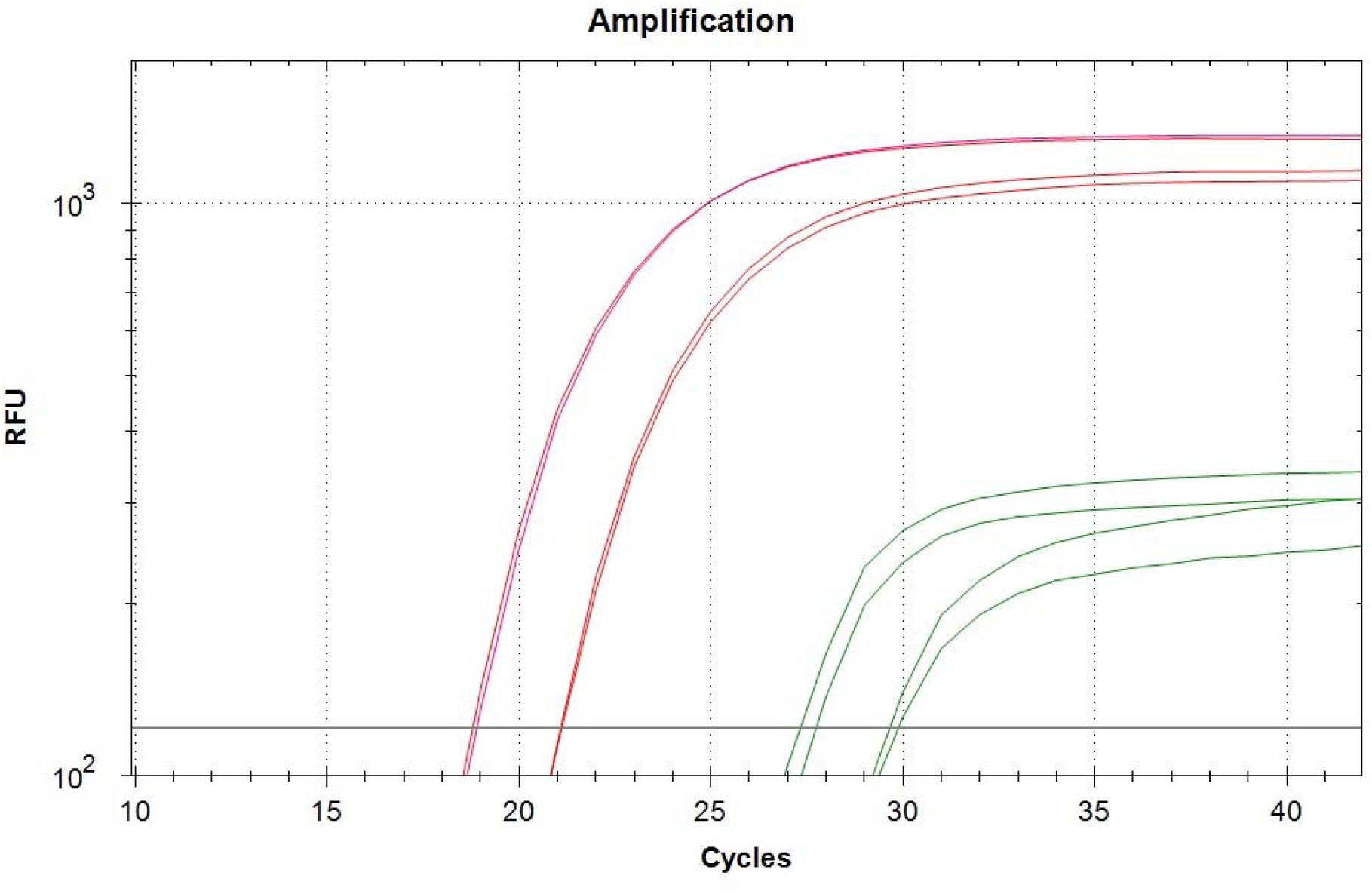
Comparison of qSDCR and qPCR assays of human B2M cDNA from two FFPE tissue blocks. The reactions contained 1 μL of the non-purified cDNA synthesized from FFPE tissue isolated RNA. The assays were performed by qSDCR (red curves) and qPCR (green curves). qSDCR significantly outperformed qPCR assays.

## 4. Discussion

The SDCR is a newly developed hybrid amplification technique which combines the advantages of isothermal amplification and PCR for NA detection, as other hybrid methods. We have compared qSDCR and qPCR techniques in the detection assays of human B2M cDNA, the bacterial gDNA of *M. bovis* and the viral gDNA of lambda bacteriophage. Selecting these objects for the tests allowed us to try out SDCR assays for different applications. B2M is a commonly used reference gene for gene expression studies and particularly in cancer research [29]. *Mycobacterium bovis* is a classical bacterial pathogen which belongs to the Mycobacterium tuberculosis complex (MBTC) and can primarily cause tuberculosis in cattle and other animals [28]. Lambda bacteriophage (*Lambdavirus lambda*) is a bacterial virus containing a 48.5 kb DNA genome.

The detection of viral and bacterial species, and particularly pathogen species, is one of the most important tasks of molecular diagnostics. We have successfully used SDCR for the assays of *Lambdavirus lambda* and *Mycobacterium bovis* gDNA (Sections 3.2 and 3.3), which provided a very high sensitivity and allowed us to detect single copies of viral and bacterial genomes in the reactions. Further, qSDCR surpassed qPCR with remarkable results in terms of amplification efficiency. The amplification factor, which shows a value of amplification per one thermocycle, exceeded 3 for the SDCR tests, versus approximately 2 for the PCR assays. As a result, the SDCR technique, in comparison with PCR, allowed for a reduction in the number of thermocycles needed for the detection of the species by a third, and shortened assay time. The use of qSDCR allowed us to detect as few as five copies of the target sequences in the reactions in fewer than 22 cycles (see Tables 1, 2), which took about 35 min. The demonstrated characteristics of the qSDCR assays, including high sensitivity and short analysis time, are very close to those previously found for other hybrid amplification techniques, such as qPCDR and q(PCR-LAMP) [18].

Another area of NA amplification is gene expression studies, which are important for biological, medical and cancer research. We have compared qSDCR and qPCR techniques in the expression analysis of a standard reference gene: human B2M. The target B2M cDNA was successfully detected by qSDCR in the library of total human cDNA and in the samples of the genetic material obtained from FFPE blocks. The materials from FFPE blocks are widely used in cancer research, but a high level of NA degradation drastically complicates the assays. Thus, these detection assays need high sensitivity and amplification efficiency. The qSDCR noticeably outperforms qPCR in the B2M tests (see Sections 3.3 and 3.4), which makes it suitable for the expression assays with complicated templates. The SDCR amplification factor was 3.32 (versus approximately 2 for PCR), and Cq values were reduced by a third in comparison with qPCR (Table 3). Thus, qSDCR appears to be a very promising application technique in gene expression studies.

Compared with other hybrid amplification methods, such as PCR-LAMP and PCDR, SDCR provides clear technological advantages in reaction architecture, primer design complexity, and enzyme utilization. LAMP and PCDR require four to six primers, which increases the risk of primer dimer formation and necessitates extensive optimization to ensure amplification efficiency. In contrast, SDCR employs only two conventional primers containing a single internal ribonucleotide, preserving the simplicity of classical PCR design while enabling strand displacement-based amplification. This streamlined primer architecture facilitates assay development and supports broad applicability across diverse nucleic acid targets.

SDCR also benefits from a simplified thermal profile. While PCR-LAMP involves a transition from an initial thermal cycling phase to an isothermal amplification stage, and PCDR requires continuous thermocycling with fine-tuned annealing conditions, SDCR uses a consistent two-step protocol with a single annealing and elongation temperature throughout the reaction. This uniform thermal regime eliminates the need for phase-specific transitions and reduces programming complexity, thereby enhancing reproducibility and assay robustness on standard real-time PCR platforms.

The usage of RNase H2 hydrolysis probes for qSDCR “real-time” amplifications provided a high assay specificity. Although the detections of the bacterial and viral gDNA were carried out in the presence of excess of human gDNA, the assays did not generate any non-specific or false-positive results. The high specificity of the qSDCRs is determined by two factors: specific probe hybridization with a target sequence and the specific hydrolysis of the probe by RNase H2.

Thermostable Pa RNase H2 would cleave the probe only in the case of a perfect matched probe– DNA duplex formation [23]. This characteristic of Pa RNase H2 suggests the possible use of qSDCR for single nucleotide polymorphism (SNP) detection.

Although SDCR employs a combination of two enzymes, SD DNA polymerase and thermostable RNase H2, the enzymatic system remains relatively simple. Both enzymatic functions, including strand displacement synthesis and site-specific nicking or probe hydrolysis, operate under the same thermal conditions and do not require stepwise enzyme addition. This simplifies the reaction workflow and reduces the potential for technical error. Moreover, the DNA polymerase used in SDCR does not require nuclease activity, as all cleavage and activation steps are mediated by RNase H2, enhancing both the specificity and modularity of the method. These features position SDCR as a technically efficient and operationally practical alternative among hybrid nucleic acid amplification strategies.

Along with the above-described advantages of SDCR over traditionally used PCR NA detection, the SDCR technique may have some limitations in use. First of all, it seems problematic to carry out a coupled reaction of reverse transcription and SDCR amplification in the “one-tube” format due to the presence of RNase H activity. We also found that SDCR exhibits the highest efficiency in the amplification of short (<200 bp) DNA fragments, so the size of the amplicons in our tests was about 100 base pairs (see Sections 2.5-2.7). However, despite possible limitations, SDCR can be a very helpful tool in NA detection, especially in diagnostics, where speed, specificity and high sensitivity are required.

Additionally, several practical aspects must be considered for broader laboratory implementation of the method. Beyond its technical merits, SDCR requires specific reagents such as SD DNA polymerase, thermostable RNase H2 and ribonucleotide-containing primers, which may increase assay costs compared to standard qPCR. For instance, SD polymerase is approximately 2.5 times more expensive than Taq polymerase. However, the markedly reduced number of cycles and shorter run time can improve throughput and partially offset these expenses, especially in high-throughput or time-critical applications.

Importantly, SDCR is fully compatible with standard real-time PCR instruments, allowing seamless integration into existing workflows without additional equipment. To support broader implementation, further studies should assess reagent availability, conduct cost-effectiveness

evaluations, and validate the method under routine diagnostic conditions. Such investigations will provide essential insights into the scalability and sustainability of SDCR for widespread clinical and research applications.

## 5. Conclusions

Upon the creation of artificial thermostable DNA polymerases with strand displacement activity, such as SD DNA polymerase, it became possible to develop new methods of NA hybrid amplification that combine the advantages of isothermal and PCR amplification. SDCR, with its high sensitivity and extraordinary efficiency of amplification, is a promising new method of hybrid amplification. We successfully applied it for the specific detection of viral and bacterial gDNA and for gene expression, including materials from FFPE blocks. A comparison between SDCR and the conventional PCR assays showed that SDCR provided an amplification efficiency more than one and a half times higher, that dramatically (at least by a third) reduced the number of thermocycles and the testing time. The described qSDCR technique is a robust, fast and very sensitive method of DNA amplification that is fully compatible with existing laboratory equipment and is suitable even for challenging samples such as FFPE. Along with other hybrid amplification techniques, SDCR could be extremely useful for NA detection and diagnostic kit development.

## Supplementary Materials

**Figure S1.**
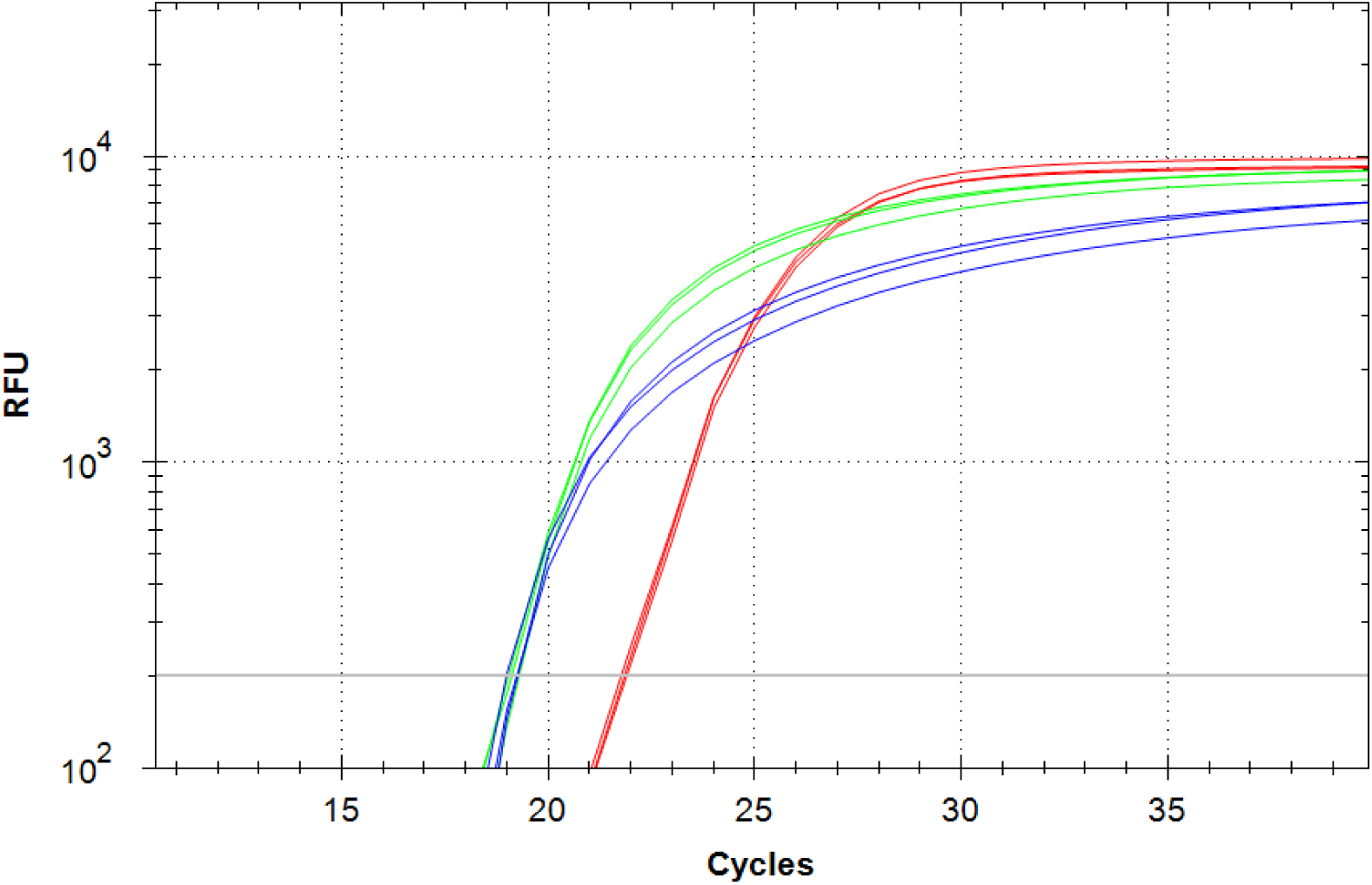
Optimization of SD polymerase concentrations for qSDCR assays. qSDCR assays of human beta-2-microglobulin (B2M) cDNA contained 0.5 pg of the human total cDNA library per reaction and different concentrations of SD DNA polymerase: 0.4 U/μL (red curves); 0.8 U/μL (green curves); 2 U/μL (blue curves). The reactions were carried out in three replicates. Concentrations over 0.8 U/µL did not improve the performance of qSDCR.

## Author Contributions

Conceptualization, K.B.I.; methodology, D.A.V., E.V.B., E.V.S., and K.B.I.; validation, V.M.K., E.V.B., K.A.B., E.V.S., and K.B.I.; formal analysis, V.M.K., K.A.B., E.V.B., E.V.S., and K.B.I.; investigation, D.A.V., E.V.B., E.V.S., and K.B.I.; resources, V.M.K. and K.B.I.; data curation, K.A.B., V.M.K., and K.B.I.; writing—original draft preparation, K.B.I.; writing—review and editing, E.V.S. and K.B.I.; supervision, K.B.I.; project administration, K.B.I.; funding acquisition, V.M.K. and K.B.I. All authors have read and agreed to the published version of the manuscript.

## Institutional Review Board Statement

Not applicable.

## Informed Consent Statement

Not applicable.

## Data Availability Statement

The original contributions presented in this study are included in the article; further inquiries can be directed to the corresponding author.

